# Niche-specific metabolic signatures in non-typeable *Haemophilus influenzae*

**DOI:** 10.64898/2026.05.26.727869

**Authors:** Keshab Bhattarai, Bikash Baral, Aleksandra Sarnowicz, Margo Diricks, Stefan Niemann, Jan Rupp, Katarzyna A. Duda

## Abstract

Non-typeable *Haemophilus influenzae* (NTHi) is a prominent opportunistic pathogen relevant to chronic respiratory diseases. NTHi’s metabolic diversity enables its survival in a wide range of environmental conditions within the host. As such, deeper research into the metabolic pathways of NTHi may open an avenue for novel therapies aimed at combating NTHi-associated respiratory diseases. Draft genome sequences from nine NTHi clinical strains from three isolation sites - ear (ear sample, ES), pharynx (pharynx sample, PS), and lower respiratory tract (Lungs) - were analyzed and annotated using RAST, PROKKA, KEGG KAAS, and antiSMASH. Pathway module coverage per-strain was computed and summarized by per-group for significant annotated metabolites. Metabolites were analyzed by LC/HRMS, identified by Metaboscape, and statistically compared using MetaboAnalyst and R software. Gene content across the tested NTHi strains was largely conserved, with limited core-genome SNP variation. Gene annotation for metabolite-related pathways revealed that all nine strains possessed largely similar sets of metabolic pathway genes, despite minor nucleotide-level differences, indicating broadly comparable metabolic capacities. In contrast, metabolomics data revealed differential metabolic profiles among the body-site groups. In a principal component analysis (PCA), the ES group was significantly separated from both the PS and Lung groups, which overlapped considerably. Detailed metabolite analyses showed that inosine, hypoxanthine, and uracil were highly significant in the ES group compared to the PS and Lung groups. For the first time, our study sheds light on the extent of metabolic differences associated with NTHi inhabiting diverse host niches. The observed metabolic differences suggest that NTHi may modulate its metabolism in a site-specific manner that is affected by environmental factors. These findings add to our understanding of how NTHi metabolism contributes to site-specific colonization.

**Author summary:** *Haemophilus influenzae* is widely recognized as a causative agent of meningitis and pneumonia. In particular, *H*. *influenzae* strains with a polysaccharide capsule—known as *H*. *influenzae* type b (Hib)—were historically a major cause of invasive disease. However, Hib has been largely eradicated following implementation of the Hib vaccine. Nonetheless, there are *H*. *influenzae* strains that lack this capsule and are therefore not targeted by the vaccine. These are known as non-typeable *H*. *influenzae* (NTHi). Following the decline of Hib, NTHi has rapidly occupied the ecological niche in the lower respiratory tract, becoming the most prominent pathogen in patients with chronic respiratory infections—particularly in those with chronic obstructive pulmonary disease (COPD), where it frequently triggers exacerbations. Importantly, NTHi is also a common component of the normal microbiome in healthy individuals, typically residing in the upper respiratory tract without causing disease.

In our study, we investigated the metabolic characteristics of NTHi isolates obtained from different body sites in patients to better understand what distinguishes strains capable of colonizing specific anatomical niches. We successfully identified several distinct metabolic features associated with NTHi strains from the ear, pharynx, and lung. These findings may serve as a foundation for future research into patient-tailored biomarkers and targeted therapies, ultimately aiming to eradicate NTHi in chronic lung infections.

## Introduction

*Haemophilus influenzae* is a human-restricted, Gram-negative pleomorphic coccobacillus that commonly colonizes the nasopharynx without any visible symptoms but can also cause disease across the respiratory tract. Strains lacking a polysaccharide capsule around the bacterial cell are termed “non-typeable” (*i.e.*, NTHi) and are now the dominant *H. influenzae* strains from mucosal infections, including acute otitis media, pneumonia, sinusitis, conjunctivitis, and exacerbations of chronic lung diseases [1]. NTHi remains a leading otitis media pathogen in children, underscoring its continuing clinical importance [2]. The rise of β-lactamase-negative ampicillin-resistant (BLNAR) strains has further complicated empirical treatment and patient care due to phenotypic resistance [3].

The first complete genome of a free-living organism was that of *H. influenzae* Rd (KW20), a landmark that initiated modern bacterial genomics [4]. NTHi lacks the capsule locus that defines serotypes a–f, explaining the “non-typeable” label and reflecting a distinct genetic strategy compared with encapsulated lineages [5].

NTHi strains are genetically diverse. They are naturally competent and strongly enriched with short “uptake signal sequences” (USS) that bias the DNA acquisition and accelerate genome remodelling [6,7]. Across hundreds of genomes, disease phenotype is only weakly predicted by core-genome multi locus genotyping phylogeny or accessory gene content, consistent with frequent microevolution and regulatory switching during niche transitions [8]. Key adhesins (HMW1/HMW2 or the trimeric autotransporter Hia) and IgA1 proteases vary among strains and are dynamically regulated during persistence, shaping adherence, invasion, biofilm development, and immune evasion in airway niches [9,10]. However, in-depth genomic potential concerning primary and secondary metabolic signatures in multiple NTHi strains has remained underexplored.

NTHi strains have adapted their metabolism to the human airway. They are auxotrophic for heme (X factor) and nicotinamide adenine dinucleotide (NAD, V factor), scavenging these cofactors from host sources through multiple outer-membrane and periplasmic systems [11]. A global transcriptomic and metabolic study reveals that NTHi strains perform “respiration-assisted fermentation,” flexibly blending fermentative pathways with electron transport to manage redox stress under microaerobic or anaerobic conditions, which are typical of mucosal biofilms and inflamed airways [12]. During long-term heme-iron restriction, NTHi strain rewire central metabolism and enhance biofilm traits that support survival [13].

Surface architecture is also metabolically encoded and niche-sensitive. The lipooligosaccharide (LOS) carries phase-variable decorations that mediate host interactions [14,15]. LOS sialylation depends on scavenged sialic acid (Neu5Ac) and associated transport/catabolic operons, conferring molecular mimicry and protection from complement, again linking metabolism to immune evasion [16]. Recent mass-spectrometry work further resolves the small-molecule and lipid collection of *H. influenzae*, revealing strain-specific phospholipid rules and previously unannotated metabolites, such as anti-microbial cyclo(Leu-Pro), but these studies rarely integrate clinical niche information [17]. Some studies have also shown substrate- (glucose, inosine, and pyruvate) and aeration-specific metabolic profiles (acetate, formate, succinate, lactate, and hypoxanthine) in two invasive-disease-related and one chronic lung disease (CLD)-related NTHi strain [12,18].

Despite extensive genomics and decades of elegant work on heme/iron acquisition, phase variation, and adhesins, we still lack a clear, systems-level map of how adaptation to clinical sites (ear, pharynx, and lower airways) imprint NTHi metabolism at the level of measurable metabolites. Large comparative genome studies show limited ability to distinguish strains from otitis media, pneumonia, CLD, and other phenotypes using sequence alone, suggesting that regulatory and metabolic plasticity, not fixed gene content, drives niche specialization.[8] NTHi strains repeatedly adapted to airway conditions (hemoglobin limitation, hypoxia, nitrosative stress) by re-prioritizing central metabolism and biofilm physiology; however, these conclusions largely derive from transcriptomics, single-strain models, or multi-isolate metabolome and niche-aware metabolomics [12,13]. We still lack comparative datasets that tie these metabolite-level features to isolation sites across diverse clinical strains [19].

We hypothesized that adaptation to different body sites corresponds to reproducible metabolic “fingerprints” that are not apparent from genome content alone. By coupling whole-genome variation to metabolite profiles, our approach aimed to reveal site-specific metabolic features that might explain persistence, inflammation, and treatment refractoriness across the NTHi disease spectrum. This niche-resolved, metabolite-centric view complements prior genomic surveys and could expose actionable metabolic vulnerabilities for diagnosis, prevention, or therapy [8,13].

## Results

### Description of the bacterial strains

Nine clinical NTHi strains previously obtained from patients diagnosed with otitis media (n=3; H104, H149, and H213) or pneumonia (n=6; H117, H158, H182, H188, H189, and H212) were included in this study [20]. Each strain was isolated from a site-specific body sample collecting ear smear (H104 and H213), bronchial secretion (H188 and H212), bronchoalveolar lavage (H117 and H158), tracheal secretion (H182) and pharyngeal smear (H149 and H189). In this study, we categorized all the strains into three major site-specific groups: ear sample (ES), pharyngeal sample (PS), and Lung. The lung group comprises all the strains isolated from the lower respiratory tract. Importantly, the strains belong to different phylogenetic clades: H-149 (PS) belongs to NTHi clade I; H-117 (Lung), H-158 (Lung) and H-212 (Lung) belong to NTHi clade IV; H-104 (ES) and H-182 (Lung) belong to NTHi clade V; H-188 (Lung) and H-189 (PS) belong to NTHi clade VIa; and H-213 (ES) belongs to NTHi clade VIb (S1 Fig and S2 Fig).

### Near-complete genomes with conserved core resistome

The genome assemblies exhibited high completeness, with BUSCO scores ranging from 98.2% to 99.4% complete single-copy orthologs, 0-0.8% fragmented, and 0-1.6% missing (S1 Table_Sheet 1), indicating near-complete genomes. Following Busco analysis, the quality of the assembly was assessed using QUAST (S1 Table_Sheet 2). QUAST assembly quality assessment indicated 29–62 contigs per genome, with total assembly lengths ranging from 1.77 to 1.98 Mb, N50 values between 77,175 and 359,254 bp, and GC contents of 37.93%–38.56% (S1 Table_Sheets 2). Gene prediction revealed between 1,750 genes in strain H104 and 2,053 genes in strain H117 (S1 Table_Sheets 5). These metrics collectively indicate high-quality genome assemblies with consistent genomic characteristics across the analysed strains.

Antimicrobial resistance genes (ARGs) analysis identified a conserved core resistome across all strains, including *hmrM* (a MATE-family efflux pump conferring resistance to acridine dyes and fluoroquinolones) and CRP (a global regulator implicated in resistance to fluoroquinolones, macrolides, and β-lactams). These genes were ubiquitous, suggesting a shared basal resistance profile (S1 Table_Sheet 3). Additionally, species-level identification via FastANI confirmed closest relatedness to *H. influenzae* NCTC 8143, with the highest ANI score of 98.86% (strain H104) (S1 Table_Sheet 4).

### Variable genome architecture

Representative sequence data for strain H188 (Fig 1a) show genomes with variable lengths and GC content (S1 Table_Sheet 5). Gene prediction revealed variability in genomic features across the analyzed\ *H. influenzae* strains. The number of protein-coding genes (CDS) ranged from 1,688 to 1,965 per genome. The genomes encoded approximately 1,800-1,900 protein-coding genes with assigned functions on average. The number of rRNA genes varied between 7 - 19, and tRNA genes ranged from 50 - 58 across strains. Additionally, all genomes contained 1 tmRNA, and one genome (H188) showed evidence of repeat regions (1-2 per genome). No pseudogenes were reported in the current annotation datasets. Core-genome SNP analysis identified pairwise SNP differences ranging from 8 to 116 SNPs among the strains, indicating moderate genomic diversity across strains. Comparative genomic visualization of NTHi strains generated using Proksee and genome-wide phylogenomic analysis of *H. influenzae* strains based on GBDP clustering are given in S2 Table and S3 Table, respectively.

**Fig 1.**
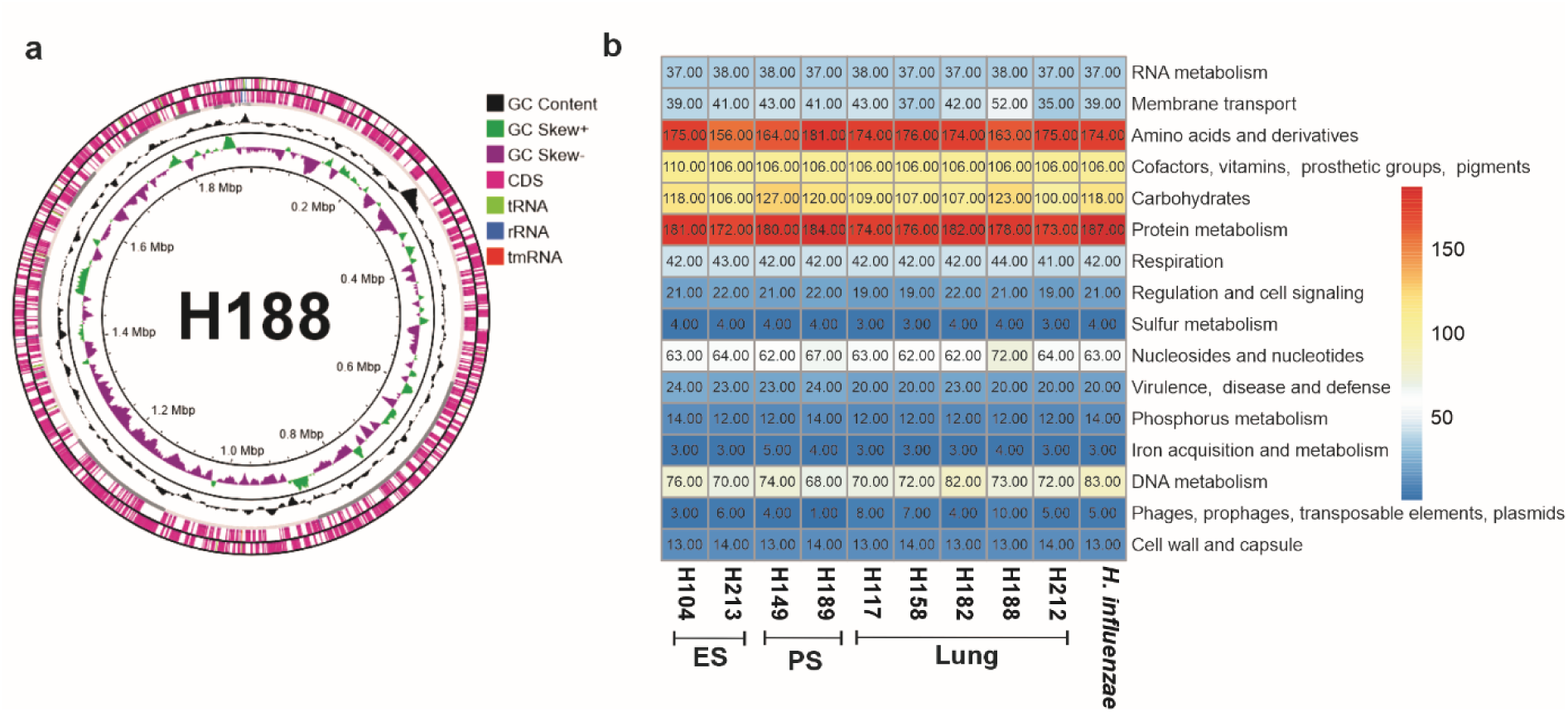
Genomic overview and accompanied metabolic extents of 9 different NTHi strains. a) Circular map and total genome information of a representative strain (H188) used in this experiment, showing the GC content, GC skew, CDS, tRNA, rRNA, and tmRNA. The circular map was created using the fasta sequence of H188 strain in Proksee (www.proksee.ca). b) Subsystem distribution of different isolated strains based on the data obtained through the RAST server (Color codes are based on their numbers obtained from the subsystems of the RAST output). Detailed data for individual cellular processes is provided in S1 Table_Sheet 6. Comparative analysis of different subsystem distributions in different isolated strains is shown. *Haemophilus influenzae* strain CP000057.2 was used as a reference.

### Conserved core functions with strain-specific metabolic variations

Based on the RAST SEED analysis, the isolated strains were found to be associated with various cellular processes, and subsystem distributions are presented as a heat-map (Fig 1b; S1 Table_Sheet 6). The RAST-based annotation revealed both conserved core functions and strain-specific variations among *Haemophilus* strains. Highly conserved subsystems across strains included RNA metabolism, cell division, fatty acid metabolism, potassium metabolism, and secondary metabolism, reflecting essential housekeeping functions. RNA metabolism was consistently represented by 37–38 genes across strains. Moderate variability was observed in amino acid metabolism (156–181 genes), suggesting differences in biosynthetic capacity and potential auxotrophies.

Strain H188 encoded a higher number of membrane transporters (52 genes), which may contribute to enhanced nutrient uptake or antibiotic resistance. Vitamin and cofactor metabolism remained largely conserved, whereas nucleotide metabolism (62–72 genes) and regulation/signaling (19–22 genes) showed mild variation that could influence stress responses and gene regulation. DNA metabolism also varied slightly; strains H182 and *H. influenzae* CP000057.2 encoded more genes (82–83), possibly reflecting enhanced DNA repair or recombination capacity (S1 Table_Sheet 6).

Virulence-associated genes ranged from 20 to 24 across strains, with H104 (ear strain) and H189 (pharynx strain) showing the highest counts. Iron acquisition genes (3–5 per genome) were present in most strains, indicating adaptation to iron-limited host environments. H188 contained the highest number of phage-related and mobile genetic elements, suggesting increased genomic plasticity and potential horizontal gene transfer. Minor variations were also observed in phosphorus, sulfur, and other metabolic pathways.

Gene-level comparisons across strains were encoded as binary presence–absence values (S1 Table_Sheets 7–9). Functional categorization based on COG classifications highlighted gene involvement in major biological processes, and the top 500 most abundant COG features were identified (S1 Table_Sheets 7–9).

### Conserved core metabolism with niche-specific adaptations

A comprehensive genomic analysis of carbohydrate metabolism pathways was conducted across the nine *Haemophilus* strains sourced from the lungs (n = 5), ear smear (n = 2), and pharyngeal smear (n = 2) as well as for the reference *H. influenzae* CP000057.2 (*H. influenzae* 86-028NP) genome. Annotation using KAAS and ortholog profiling revealed a broadly conserved core set of metabolic genes underpinning central carbon metabolism, alongside targeted, niche-correlated gene losses indicative of differential metabolic adaptation (Fig 2) (S4 Table_Sheets 1, 2, and 3).

**Fig 2.**
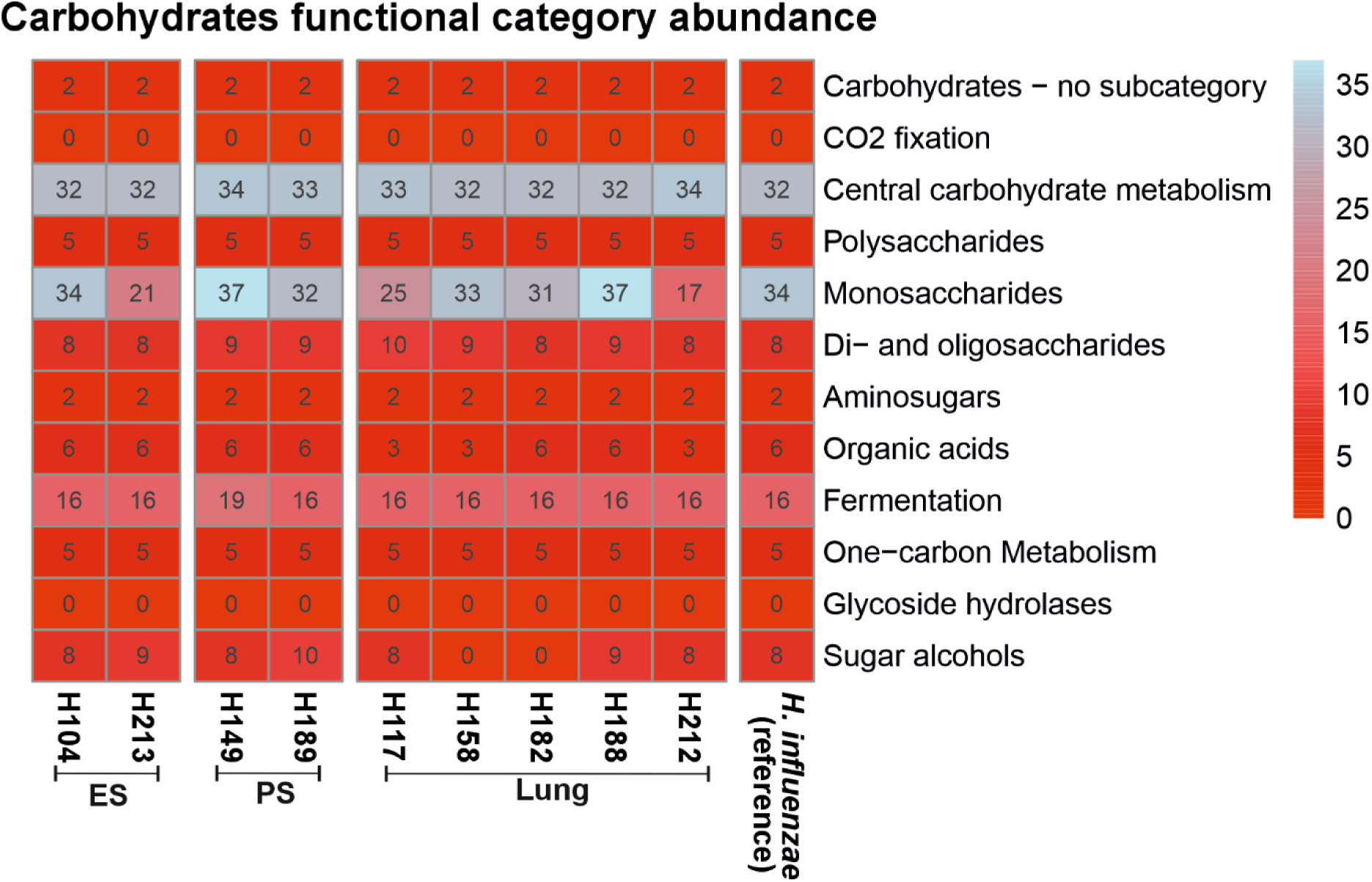
Carbohydrate metabolism encoding genes found in different studied *Haemophilus influenzae* strains. The numbers represent the number of KO (KEGG ORTHOLOGY) entries retrieved by KAAS from the KEGG database.

Among 19 targeted metabolic genes, 17 were universally conserved across the strains, including *pfkA*, *pyk*, *pckA*, and the fructose-1,6-bisphosphatase isoforms (*fbp*, *glpX*). The only exception was *frmA*, which was absent in the reference strain. Notably, the ear strains (H104, H213) retained the complete set. While this may suggest metabolic robustness within this niche, the limited number of ear strains warrants caution, and broader sampling would be required to confirm whether this pattern reflects niche-specific metabolic stability or simply isolate-level variation. Pyruvate metabolism (26 genes) showed near-complete conservation, with *frmA* again missing in the reference strain. Likewise, starch and sucrose metabolism genes (*glgA*, *glgB*, *galU*, *pgi*, *glgC*, *malQ*, *crr*) were conserved, although glycogen phosphorylase (*glgP*) was universally absent, suggesting divergence in carbohydrate storage as compared to the reference strain. The Leloir pathway (*galK*, *galT*, *galM*, *galU/UGP2*) was also intact, but *galE* was consistently undetected, possibly reflecting annotation gaps or alternative mechanisms of galactose-1-phosphate conversion into UDP-galactose (S4 Table_Sheets 1, 2 and 3).

In contrast, the tricarboxylic acid (TCA) cycle and related pathways showed localized variation. Of 13 TCA genes, 12 were conserved, including *mdh*, *aceE*, *pdhC*, *lpd*, and *frdA–D*. The β-subunit *sucC* was absent only in lung strain H182, while its α-subunit (*sucD*) and other components remained intact. Similarly, propanoate and butanoate pathways were broadly conserved, with *ackA*, *pta*, *accA–D*, *mgsA*, *frdA–D*, *atoA*, *ilvH*, and *ilvN* present. However, *sucC* was again missing in H182, *pta* in H149, and *pflD* was universally absent. Large acetolactate synthase subunits (*ilvB*, *ilvG*, *ilvI*) appeared sporadically, mainly in PS strain H149. Glyoxylate and dicarboxylate metabolism was largely conserved (*mdh*, *glyA*, *ACAT*, *glxK*, *gph*, *purU*, *eda*, *glnA*), though lung strains showed selective absences (*ACAT*, *glxK*, *eda* in H117; *glxK* in H212). By contrast, inositol phosphate metabolism was fully conserved across all strains.

Sugar interconversion and accessory pathways exhibited greater variability. Within the pentose phosphate pathway (PPP), 14 of 17 genes were conserved, but *kdgK* and *eda* were absent in lung strain H212, while *fbp* was missing in pharyngeal strain H189. Pentose and glucuronate interconversion genes consistently included *galU*, *RPE*, *fucK* and *fucI*, whereas other genes like *xylA*, *xylB*, *fucA*, *uxuA* and *uxaC* were selectively absent, particularly in H212 (S4 Table_sheet 2 and 3).

Fructose and mannose metabolism was mostly conserved (*fbaA*, *tpiA*, *fruA*, *fbp*), though *fucA* genes were absent in H213, and annotation gaps were noted for *fruK* and *manB*. Finally, C5-branched dibasic acid metabolism was largely conserved across all strains, with key enzymes such as *leuB*, *leuC*, *leuD*, *ilvH*, *ilvN*, and *sucD* present in every genome examined. However, notable strain-specific variation was observed in the acetolactate synthase large subunits (*ilvB*, *ilvG*, *ilvI*), which were exclusively detected in strain H149 (S4 Table_sheets 2 and 3). This suggests that while the core pathway remains broadly intact, the capacity for certain branched-chain amino acid biosynthetic reactions may be uniquely maintained in H149. Collectively, these results highlight the strong conservation of central metabolism, alongside selective gene losses indicative of niche-specific remodelling and functional redundancy.

### Deprived biosynthetic potential of secondary metabolites

Since secondary metabolites play ecological roles in competition, defence and communication, we first checked the genomic capacity for producing secondary metabolites in the tested NTHi strains using antiSMASH analysis of the whole genome sequences. Surprisingly, the analysis revealed only one secondary metabolite biosynthetic gene cluster, encoding an azole-containing ribosomal peptide, which was conserved in all strains (S5 Table). This conserved gene cluster showed no similarity to any known biosynthetic gene clusters or ribosomal peptides.

### Metabolomic profiling revealed diverse primary metabolites

After cultivating each of nine NTHi strains in chemically defined liquid medium, the resulting cell-free supernatants were subjected to metabolite identification. This setup allowed us to characterize both substrate consumption and metabolite release under controlled conditions, providing insight into metabolic activities potentially relevant to different host niches.

MetaboScape analysis initially showed 221 features altogether from 48 samples in positive MS polarity. Seventy-five features were detected as artefacts (eluted before 0.5 min, lacking MS^2^ spectra and present extensively in n-butanol blank samples) and excluded. Out of the remaining 146 features, only 38 features were found to be produced or consumed by at least one of the nine NTHi strains (S6 Table), while the rest were predominant artefacts from background noise and methanol blanks, and hence excluded.

Out of these, 18 features were successfully annotated against MS^2^-spectral library with MS/MS score above 900 (S3 Fig), where inosine, hypoxanthine, niacinamide (also known as nicotinamide), uracil and isoleucine are annotated for multiple features. Inosine is detected two times with *m/z* 269.08850 [M + H]^+^ at 1.20 min and *m/z* 269.08863 [M + H]^+^ at 1.77 min while hypoxanthine is detected three times with *m/z* 137.04562 [M + H]^+^ at 1.20 min, *m/z* 137.04561 [M + H]^+^ at 1.76 min and *m/z* 137.04571 [M + H]^+^ at 1.61 min. Niacinamide is detected three times with *m/z* 123.05522 [M + H]^+^ at 1.20 min, *m/z* 123.05523 [M + H]^+^ at 1.52 min and *m/z* 123.05510 [M + H]^+^ at 1.76 min. Uracil is detected three times with *m/z* 113.03432 [M + H]^+^ at 1.22 min, *m/z* 113.03432 [M + H]^+^ at 1.58 min and *m/z* 113.03433 [M + H]^+^ at 1.75 min. Isoleucine is detected twice with *m/z* 132.10187 [M + H]^+^ at 1.20 min and *m/z* 132.10193 [M + H]^+^ at 1.77 min. These repeatedly annotated features are either differed by retention time or *m/z* value or both above the tolerance level but have identical MS^2^ spectra. The other annotated metabolites are detected as phenolsulfonphthalein (also known as phenol red) with *m/z* 355.06363 [M + H]^+^ at 6.97 min, phenylalanine with *m/z* 166.08643 [M + H]^+^ at 2.24 min, pyridoxine with *m/z* 170.08153 [M + H]^+^ at 1.17 min, valine with *m/z* 118.08616 [M + H]^+^ at 1.17 min and arginine with *m/z* 175.11932 [M + H]^+^ at 1.06 min. Phenolsulfonphthalein is found to be an acidic product of hydrolysis of sodium salt of phenol red (molecular weight: 376.4 Dalton) present in the culture medium. The unannotated features are denoted as “retention time_*m/z*”. Eight of annotated metabolites (Fig 3) were significantly asymmetrically distributed among 9 NTHi strains (see section 3.8).

**Fig 3.**
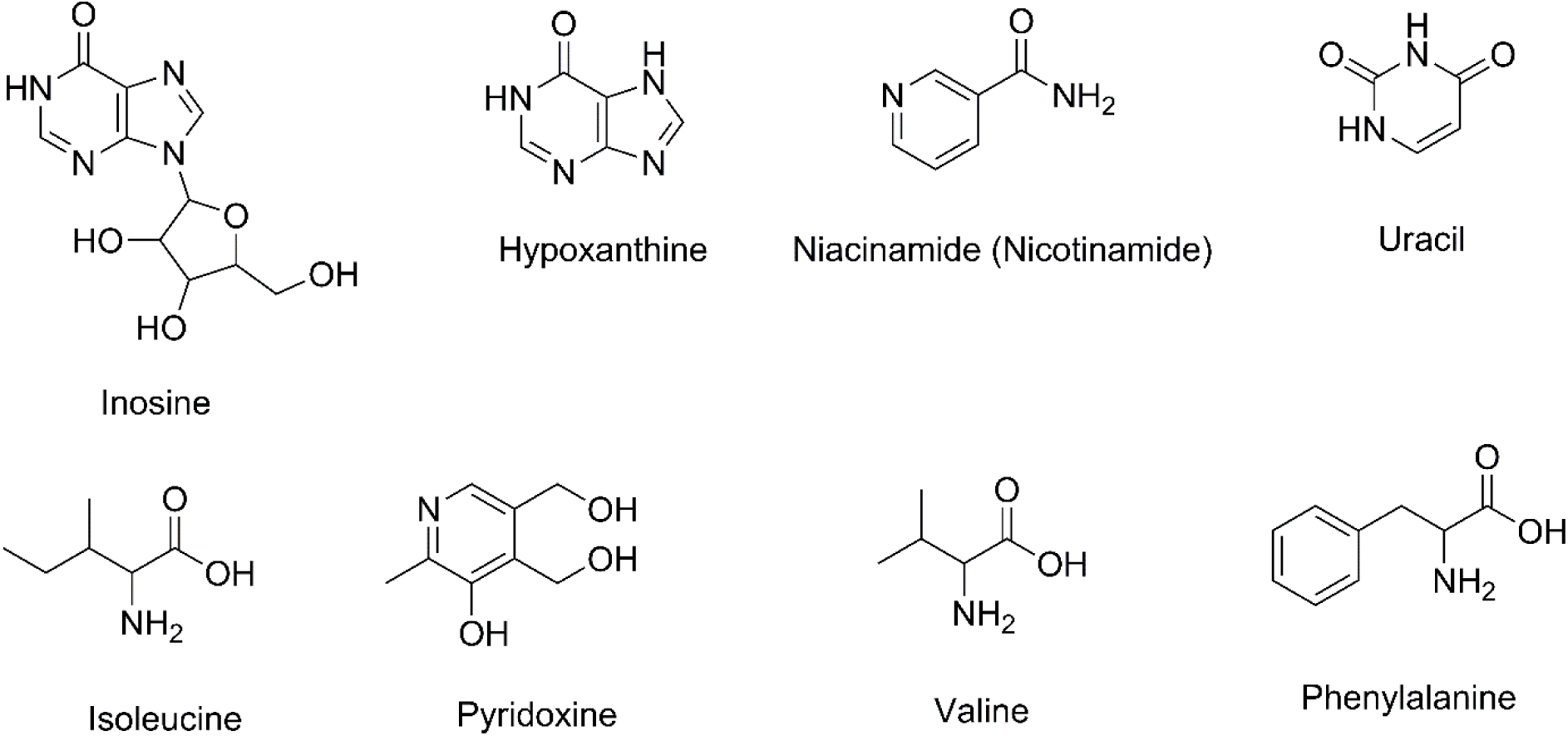
Chemical structures of metabolic signatures from site specific samples.

### Statistically distinct metabolic signatures by isolation site

Thirty-eight samples (from 9 NTHi strains and media control) and 38 features were analysed through interactive principal component analysis and linear modelling using MetaboAnalyst 6.0. A synchronized 3D PCA plot (PC1: 44.4%, PC2: 15.0%, PC3: 10.3%) showed that ES samples are separated to the upper left from the PS and Lung samples while visualizing on metadata-overlay (S4 Fig). A 2D score plot with 44.4% PC1 (x-axis), 15% PC2 (y-axis), F-value: 10.82, R-squared: 0.48841 and *p*-value (based on 999 permutations): 0.001 showed that ES still edged to the upper left region as compared to overlapped PS and Lung groups while visualizing on 95% confidence regions (Fig 4).

**Fig 4.**
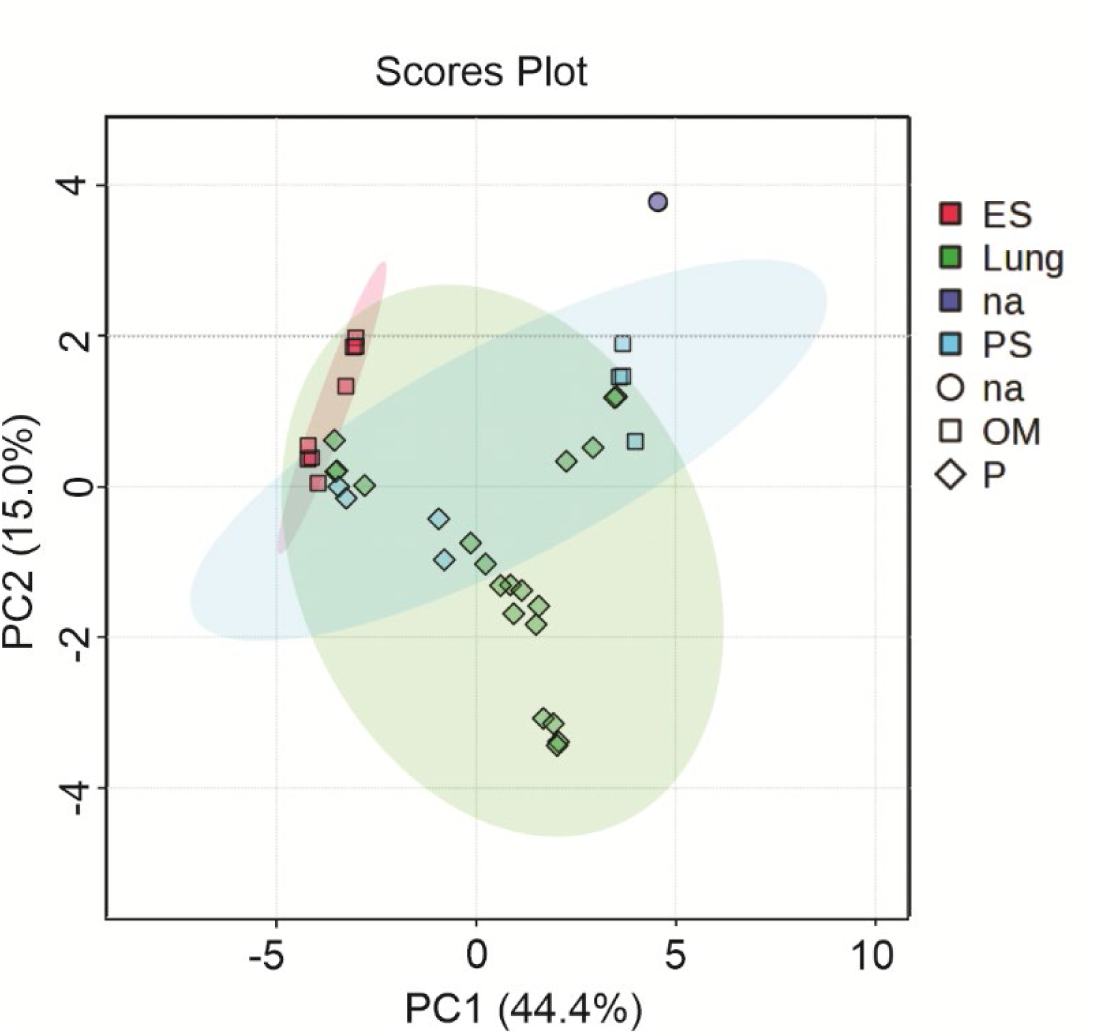
PCA score plot of isolation site-specific samples (PC1 vs. PC2) based on metabolite intensities. (ES = ear samples, PS = pharyngeal sample, OM = otitis media, P = pneumonia, na = medium control).

The linear models with *P*-value cut-off: 0.05 (False Discovery Rate) have pointed out a total of 28 significant metabolites varied in site-specific groups: ES, PS, and Lung. The model shows 23, 22, and 2 significant metabolites differed in ES vs Lung, ES vs PS, and Lung vs PS groups, respectively. The list of significant metabolites varied in different groups with their respective logFC (fold change) and P-values are given in S7 Table(a-c). Out of these metabolites, 14 are annotated against MS^2^-spectral libraries with repeated annotation of inosine (2 times), hypoxanthine (3 times), uracil (2 times) and niacinamide (3 times) along with, phenylalanine, isoleucine, pyridoxin and valine. All significant metabolites after merging the repeated annotations and their abundance ranking in different sample groups are given in Table 1. The most significantly differed metabolite in ES vs Lung and ES vs PS is the unknown compound “1.45_136.06189”. This metabolite along with “1.15_136.06189” gives a closer hit with adenine while annotating with reduced threshold of MS/MS score (700-800). Additionally, inosine, hypoxanthine, niacinamide and uracil are major significantly varied metabolites in both ES vs Lung and ES vs PS. Phenylalanine and isoleucine are only significantly differed metabolites in ES vs Lung, while pyridoxin and valine are only significantly differed metabolites in ES vs PS. Besides, phenylalanine and an unknown metabolite, “1.13_104.10697”, are the only compounds that significantly differed in the Lung vs PS groups.

**Table 1.**
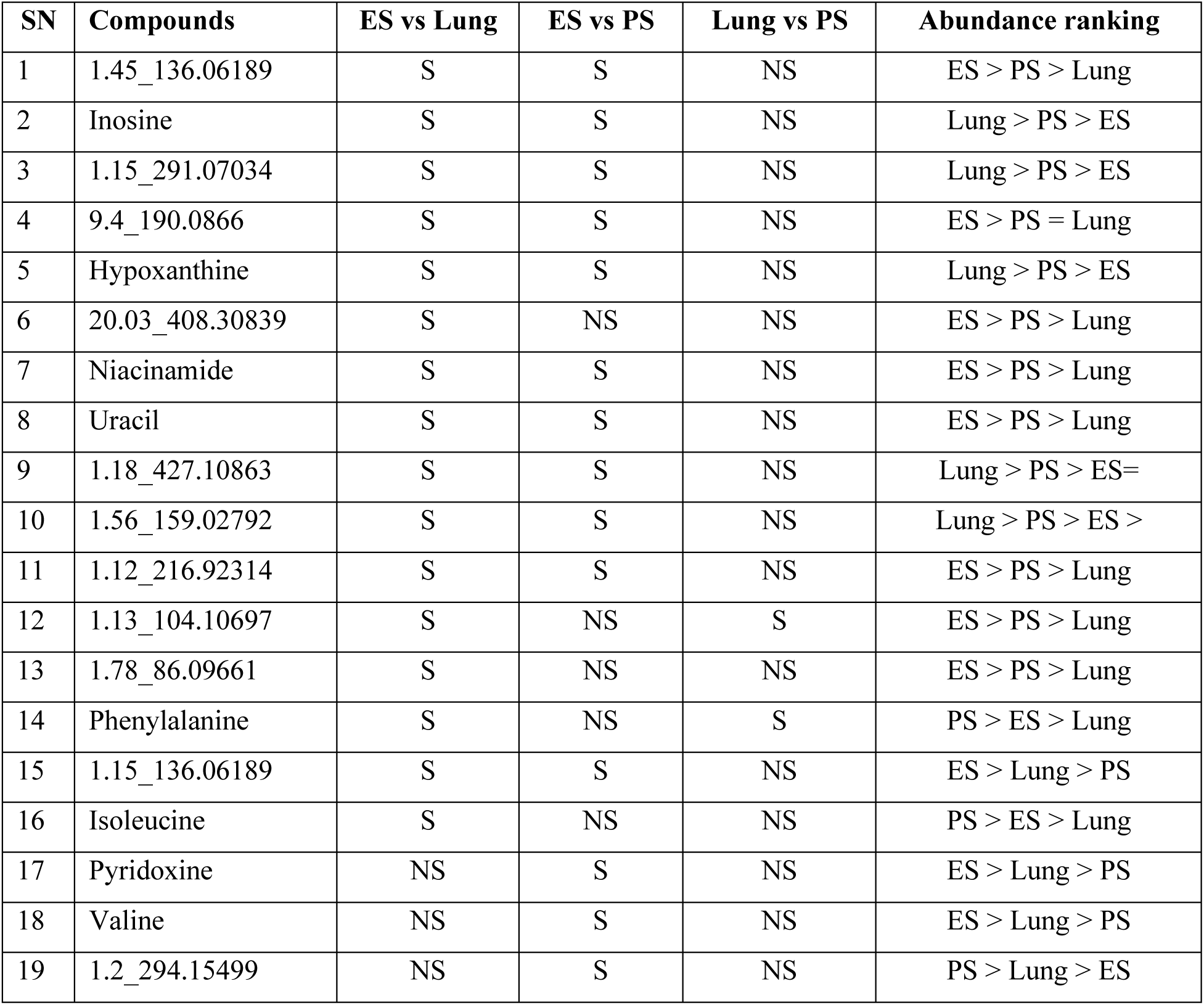
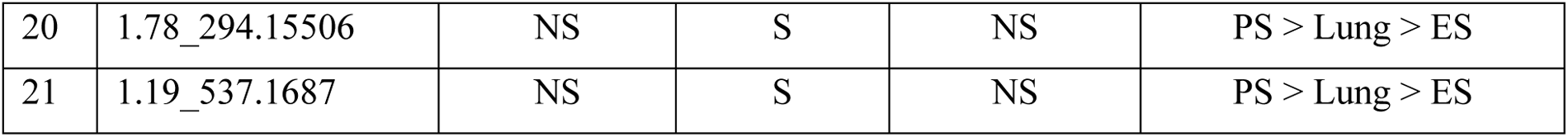
Summary of significantly produced or consumed metabolites in ES vs Lung, ES vs PS, and Lung vs PS ranked by abundance. (S = significant, NS = non-significant, ES = ear sample, PS = pharyngeal sample).

Box plots of significant metabolites generated using R (tidyverse package) after merging the repeatedly annotated metabolites by average, shows the extent of abundance in different site-specific groups (Fig 5).

**Fig 5.**
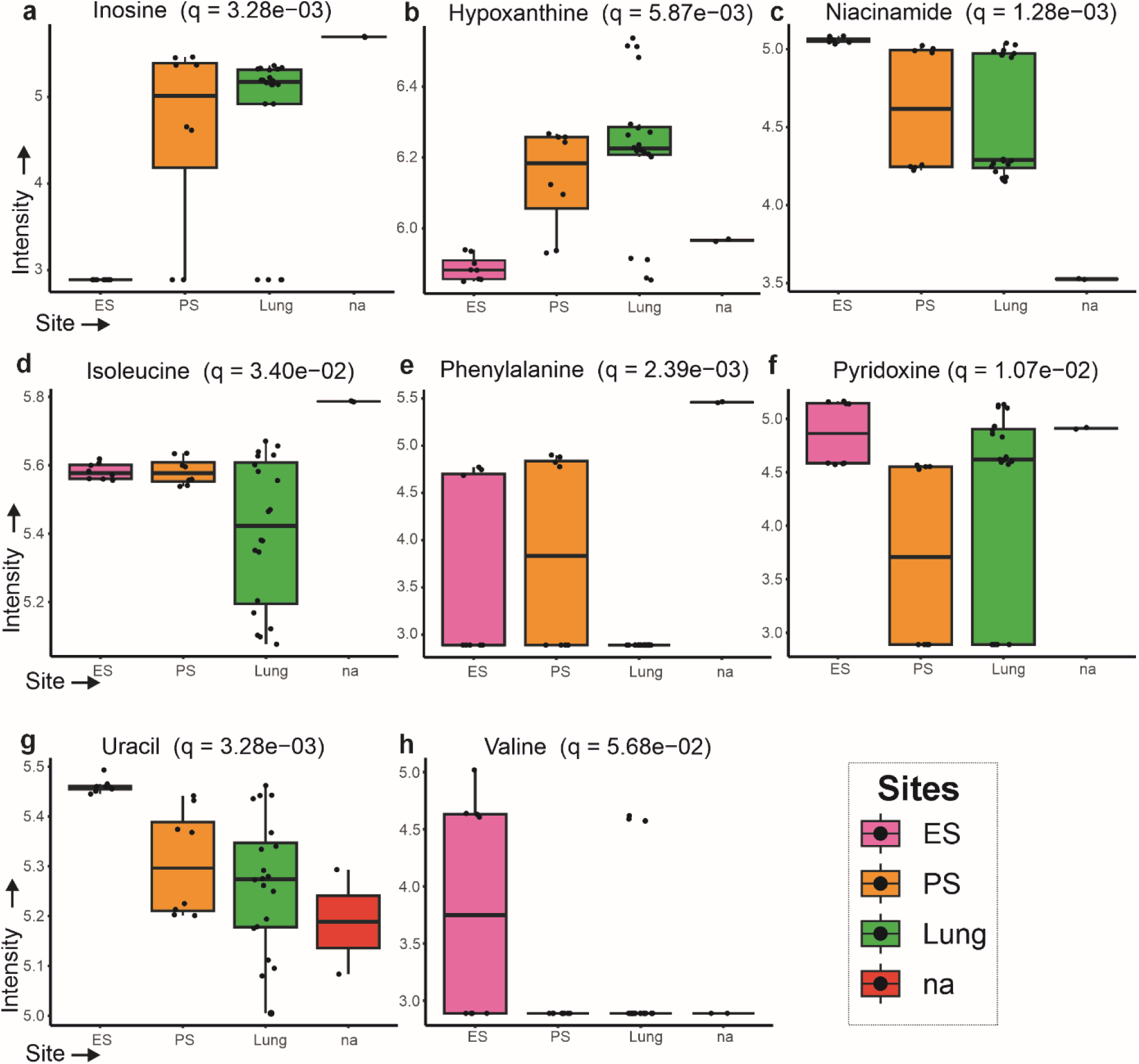
Box plots of significantly varied metabolites in site-specific samples. The *q* values presented in each box plot represent the Kruskal-Wallis false discovery rates, FDR. (na = medium control, ES = ear sample, PS = pharyngeal sample).

The samples of the strains from ES contained lower amount of hypoxanthine than that from PS and Lung and no inosine. The samples of the ES group contained higher amount of niacinamide, uracil, pyridoxine than that of PS and Lung. ES strains very also the only one showing the presence of valine. Samples from PS group contained the higher amount of phenylalanine as compare to ES while Lung group contained no phenylalanine. Samples from Lung group contained lower amount of isoleucine than that from ES and PS.

### Uniformly conserved genes despite of heterogenic metabolome

In the KAAS analysis, the genes for all metabolic pathways of the significantly different metabolites between the strains from different isolation sites were found in all 9 NTHi strains. These were genes related to purine metabolism, nicotinate and nicotinamide metabolism, pyrimidine metabolism, phenylalanine biosynthesis, phenylalanine metabolism, valine-leucine-isoleucine biosynthesis, valine-leucine-isoleucine degradation, and vitamin B6 metabolism. Extracted KEGG orthology (KO) entries which resemble a defined enzyme activity, protein complex, or functional subunit accompanied by each site-specific group (unified from individual strains) per pathway showed more than 90% identical distribution for purine metabolism, pyrimidine metabolism, and nicotinate and nicotinamide metabolism, while 100% identical distribution for the rest of the pathways (Table 2).

**Table 2.**
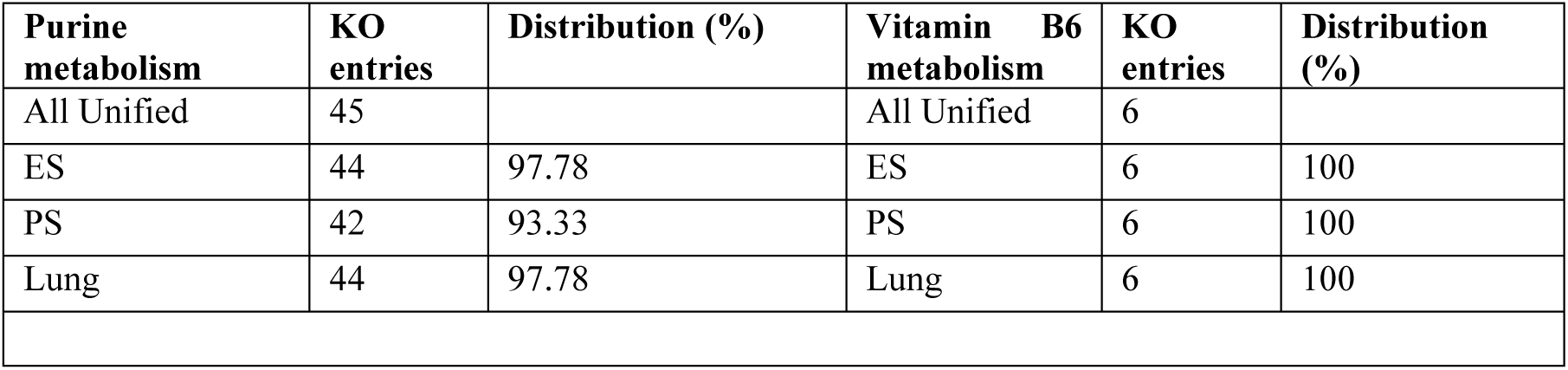
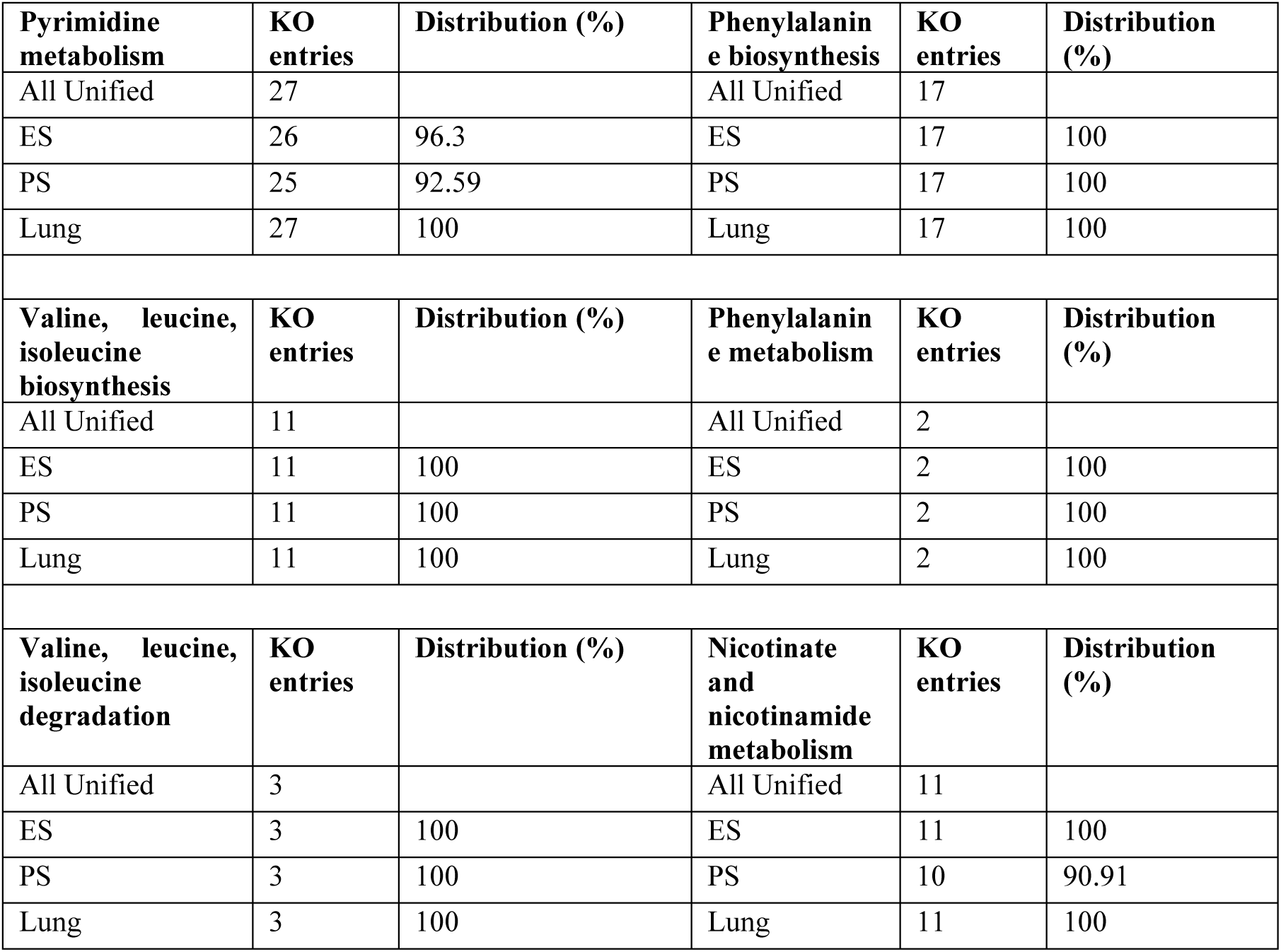
Retrieved number of KO entries of NTHi strains by group for selected KEGG pathways. (ES = ear sample, PS = pharyngeal sample).

| Purine metabolism | KO entries | Distribution (%) | Vitamin B6 metabolism | KO entries | Distribution (%) |
| --- | --- | --- | --- | --- | --- |
| All Unified | 45 |  | All Unified | 6 |  |
| ES | 44 | 97.78 | ES | 6 | 100 |
| PS | 42 | 93.33 | PS | 6 | 100 |
| Lung | 44 | 97.78 | Lung | 6 | 100 |

| **Pyrimidine metabolism** | **KO entries** | **Distribution (%)** | **Phenylalanine biosynthesis** | **KO entries** | **Distribution (%)** |
| --- | --- | --- | --- | --- | --- |
| All Unified | 27 |  | All Unified | 17 |  |
| ES | 26 | 96.3 | ES | 17 | 100 |
| PS | 25 | 92.59 | PS | 17 | 100 |
| Lung | 27 | 100 | Lung | 17 | 100 |
| **Valine, leucine, isoleucine biosynthesis** | **KO entries** | **Distribution (%)** | **Phenylalanine metabolism** | **KO entries** | **Distribution (%)** |
| All Unified | 11 |  | All Unified | 2 |  |
| ES | 11 | 100 | ES | 2 | 100 |
| PS | 11 | 100 | PS | 2 | 100 |
| Lung | 11 | 100 | Lung | 2 | 100 |
| **Valine, leucine, isoleucine degradation** | **KO entries** | **Distribution (%)** | **Nicotinate and nicotinamide metabolism** | **KO entries** | **Distribution (%)** |
| All Unified | 3 |  | All Unified | 11 |  |
| ES | 3 | 100 | ES | 11 | 100 |
| PS | 3 | 100 | PS | 10 | 90.91 |
| Lung | 3 | 100 | Lung | 11 | 100 |

In purine metabolism, we observed three genes *deoD* (K03784), *LACC1* (K05810), and *ushA* (K11751) catalyzing inosine interconversion to hypoxanthine, inosine synthesis from adenosine, and inosine synthesis from inosine monophosphate (IMP), respectively. Additionally, we detected two genes directly involved with hypoxanthine: *hprT* (K00760), which catalyzes hypoxanthine to IMP, and again *deoD* (K03784), which catalyzes the interconversion of hypoxanthine and deoxyinosine. This indicates that all the strains have equivalent genomic potential for the metabolism of inosine and hypoxanthine.

In the nicotinate and nicotinamide metabolism pathway, we detected a gene *cobB* (K12410), responsible for catalyzing NAD^+^ directly to niacinamide. Furthermore, gene *nadR* (K06211) and gene *deoD* (K03784) in combination can catalyze NAD^+^ to niacinamide through intermediate steps.

In pyrimidine metabolism pathway, we detected genes *udp* (K00757) that interconverts uridine and deoxyuridine to uracil. A*spC* (K00813) and *hisC* (K00817), genes of the phenylalanine metabolism pathway were present in all 9 strains. They can independently interconvert L-phenylalanine to phenylpyruvate (L-Phenylalanine + 2-Oxoglutarate <=> Phenylpyruvate + L-Glutamate).

In valine, leucine and isoleucine degradation pathway, we detected a gene, E2.6.1.42 (K00826), that catalyzes the L-isoleucine interconversion to (S)-3-Methyl-2-oxopentanoic acid and L-valine interconversion to 3-methyl-2-oxobutanoate. This gene was present in all 9 strains. We also detected a complete set of genes, E2.2.1.6L (K01652), E2.2.1.6S (K01653), *ilvC* (K00053), *ilvD* (K01687), E2.6.1.42 (K00826), and *alaA* (K14260), that catalyze pyruvate to L-valine and L-isoleucine through multiple intermediate steps.

In vitamin B6 metabolism, we detected a gene *pdxK* (K00868) that catalyzes pyridoxine to pyridoxine-5’ phosphate and another gene *pdxH* (K00275) that catalyzes the interconversion of pyridoxine to pyridoxal and further to pyridoxamine.

In purine metabolism among all three site-specific groups, K20881 (*yrfG*, IMP to Inosine) was unique to ES, K25589 (*inuH*, inosine to hypoxanthine, uridine to uracil) was unique to Lung and K01081 (E3-1-3-5, IMP to Inosine, AMP to adenosine, UMP to uridine) was unique to ES and Lung while the rest were present in all groups. In nicotinate and nicotinamide metabolism, K01081 was unique to ES and Lung. In pyrimidine metabolism, K25589 was unique to Lung, while K01081 was unique to ES and Lung. Distribution of KO entries per strain is given in S8 Table.

## Discussion

NTHi strains are clinically significant opportunistic pathogens, particularly in vulnerable patients with respiratory tract infections. Due to the absence of capsular polysaccharides, NTHi strains are not targeted by the Hib vaccine and have emerged as dominant agents in invasive *H*. *influenzae* disease. Similar to other bacterial pathogens, NTHi strains are thought to possess a versatile metabolic repertoire that enables adaptation to diverse host microenvironments. We hypothesized that NTHi strains adapted to different anatomical sites (ear, pharynx, and lower respiratory tract) might exhibit distinct metabolic profiles, reflecting niche-specific substrate utilization and metabolite production. To investigate this, we integrated whole-genome sequencing with LC/MS-based metabolomics to assess whether metabolic phenotypes could be inferred directly from genomic data.

High-quality genome assemblies of the strains isolated from three different body sites, supported by BUSCO completeness scores (98.2-99.4%) and robust QUAST metrics, confirmed their suitability for downstream genomic analyses [21–23].

Our pangenome and functional subsystem analyses demonstrated a highly conserved core genome, particularly in essential functions such as RNA metabolism, fatty acid biosynthesis, and cell division [24,25]. In contrast, we observed variation in amino acid and carbohydrate metabolism, as well as membrane transport, suggesting niche-specific adaptations [26]. Notably, strains H149 and H188 isolated from PS and bronchial secretion (site Lung) respectively, exhibited enhanced glycan degradation capacity, likely facilitating colonization of mucosal surfaces [27–30]. Strain H188 isolated from Lung displayed increased mobile elements, suggesting greater genomic plasticity and potential for horizontal gene transfer [31–37]. Our comparative genomic analysis highlights the conservation of a carbohydrate metabolic framework across *H. influenzae* strains investigated from diverse human anatomical sites, emphasizing its crucial role in bacterial survival, colonization, and pathogenesis [12,38]. The core glycolytic, gluconeogenic, and TCA cycle genes showed strong evolutionary conservation, essential for energy production and biosynthesis under variable host conditions [39,40]. Interestingly, some lung and pharyngeal strains exhibited selective loss of alcohol dehydrogenase (*frmA*) and fructose-1,6-bisphosphatase II (*glpX*), likely reflecting adaptive metabolic modulation in response to niche-specific substrate availability and functional redundancy [41,42]. Pentose phosphate pathway genes critical for NADPH generation remain largely conserved, while targeted losses of ribokinase, Entner-Doudoroff components, and gluconeogenic regulators indicate metabolic streamlining in lung and pharyngeal niches [43].

Variability in pentose, glucuronate, fructose, mannose, and fucose metabolism genes, particularly losses in some ear and lung strains, points to selective suppression of peripheral carbohydrate pathways, probably driven by nutrient limitations or host immune pressures [44,45]. The retention of galactose metabolism genes essential for mucosal colonization contrasts with an undetected *galE*, suggesting annotation gaps or alternative enzymatic routes [46]. The starch and sucrose pathway shows preserved glycogen biosynthesis with universal absence of glycogen phosphorylase, hinting at a strategic shift favoring carbohydrate storage over rapid degradation [47].

Pyruvate metabolism genes are highly conserved, except for lactate dehydrogenases and acetyl-CoA metabolism genes lost in ES strains, consistent with localized metabolic contraction [48–50]. Similar losses of genes involved in glyoxylate, propanoate, and butanoate pathways reinforce this niche-driven metabolic genome reduction, likely adapting to ear-specific environmental constraints [12,51]. Notably, the restricted presence of acetolactate synthase large subunits in ES strains and unique retention of succinyl-CoA synthetase β-subunit in Lung strains highlight subtle strain- and niche-specific adaptations impacting amino acid and TCA metabolism [12,52]. In contrast, inositol phosphate metabolism genes exhibit invariant conservation, reflecting their indispensable role in membrane phospholipid synthesis and bacterial signaling across niches [53–56].

We then shifted our analysis to focus on secondary metabolism, and *in silico* results revealed that all tested NTHi strains lacked the biosynthetic potential to produce secondary metabolites except one unknown ribosomal peptide. Secondary metabolites mainly play ecological roles such as competition from other microbes, survival, defence and microbial communication. Lacking biosynthetic gene clusters for diverse secondary metabolites may indicate their evolution in an ecological niche that doesn’t explicitly require such functions [57,58]. This means that all metabolites detected in the LC-MS analysis were primary metabolites, which are physiologically essential for cellular division and growth. Notably, the samples from Lung and PS contained higher amounts of inosine and hypoxanthine compared to ES. Since inosine was a part of the culture medium, it appears that ES strains consumed it completely. Inosine can only be metabolized to hypoxanthine, which then acts as an intermediate in the purine degradation process leading to the formation of metabolites such as xanthine and urea. As part of parallel reactions occurring in this process, related purine bases such as adenine and guanine could also be formed. It means that the metabolism of inosine in PS and Lung strains stops at the level of hypoxanthine, but in ES strains hypoxanthine is further metabolised.

Inosine and hypoxanthine are crucial metabolic intermediates in purine metabolism, which is also a dominant pathway in bacterial metabolism. In humans, inosine has been shown to have anti-inflammatory effect in allergic lung inflammation and regulates cytokine generation. In the airways, inosine exerts protective effects, reducing the levels of pro-inflammatory cytokines such as tumour necrosis factor-α, interleukin-1β, interleukin-6, and macrophage inflammatory protein-2, reducing leukocyte recruitment into lung tissue [59,60]. Hypoxanthine is a significant biomarker in COPD patients, with elevated levels in serum compared to healthy controls [61]. Elevated hypoxanthine and other adenyl-purine products have also been observed in airway BAL fluid of inflamed lungs, particularly in cystic fibrosis preschoolers with neutrophil inflammation [62,63]. Across human BAL fluid and animal pneumonia models, purine catabolites (inosine and hypoxanthine), NAD-axis metabolites and free amino acids strongly track neutrophilic inflammation and structural lung disease [62–64]. Interestingly, *in vivo* RNA-seq analysis of lungs has revealed up-regulation of NTHi *de novo* purine biosynthesis and fitness dependence as compared to wild type NTHi strains during murine lung infection model. The release of purine breakdown products by PS and Lung strains is consistent with substantial purine salvage in airway-adapted strains, contributing to inflammation [65].

Niacinamide, a primary precursor of NAD^+^, plays a crucial role in nicotinate and nicotinamide metabolism. Notably, all strains produced niacinamide, but the ES group showed significantly higher levels. Functionally, niacinamide is involved in energy processes, DNA repair, redox reactions, and cellular stress mechanisms. Previous studies have demonstrated that niacinamide acts as an antimicrobial compound against bacteria and fungi by arresting the microbial cell cycle [66]. Furthermore, elevated niacinamide and NAD^+^ levels have been observed in BAL fluid from early cystic fibrosis patients during neutrophilic inflammation [63]. Niacinamide has also been shown to suppress lung injury and proinflammatory mediator accumulation by inhibiting MAPK and AKT/NF-κB signaling pathways [67]. *H. influenzae* is unable to synthesize NAD and relies on NAD/nicotinamide riboside/nicotinamide mononucleotide salvage, the detection of niacinamide in the supernatant is compatible with NAD turnover by these pathways [68–71]. However, the site-specific relevance of niacinamide in the human body remains unknown and warrants further investigation.

Pyridoxine (vitamin B6) was utilized by all strains, but the PS strains showed the highest utilization compared to ES and Lung strains. In bacteria, pyridoxine plays a crucial role as a coenzyme in various reactions, primarily involved in amino acid metabolism. Additionally, pyridoxine exhibits potent antioxidant activity, effectively neutralizing reactive oxygen species and contributing to cellular well-being [72]. In some pathogenic bacteria such as *Streptococcus pneumoniae*, *Helicobacter pylori* and *Mycobacterium tuberculosis*, vitamin B6 biosynthesis is very essential for its survival, motility and virulence [73].

Uracil, a product of pathogenic bacteria, serves as a mucosal signal that triggers host dual oxidase (DUOX) to produce antimicrobial reactive oxygen species (ROS). This leads to increased extracellular uracil and enhanced microbe-immune cross-talk [62,74–76]. During nucleoside catabolism of uridine, uracil and ribose are generated. Notably, genes involved in nucleotide metabolism are more commonly found in pathogens than commensal bacteria. Furthermore, ribose, a by-product of nucleoside catabolism, induces bacterial quorum sensing (QS) and virulence gene expression, which can trigger a commensal-to-pathogen transition [77]. In our study, we observed that uracil was produced in the highest amounts in ES samples compared to PS and Lung samples.

Phenylalanine, an essential aromatic amino acid, is produced by many bacteria and is one of the nine essential amino acids in the human diet [78]. In inflamed BAL fluid of cystic fibrosis patient free amino acids including phenylalanine, valine and isoleucine increase with neutrophil count (host proteolysis) [63]. Furthermore, elevated phenylalanine levels in lung and blood samples have been shown to induce inflammation by enhancing the innate immune response and releasing proinflammatory cytokines. This can also promote pyroptosis of alveolar macrophages, exacerbating lung inflammation and acute respiratory distress syndrome lethality [79]. However, in contrast to published data, our study found that all tested strains metabolised phenylalanine, with the majority of strains isolated from Lung.

Biosynthesis of branched-chain amino acids, such as valine, leucine, and isoleucine, is crucial for bacterial survival and growth [80]. The study of Begona et al. found that NTHi up-regulates non-aromatic amino acid (leucine, isoleucine, and valine) biosynthesis during lung infection [65]. Furthermore, research suggests that a decrease in valine can increase competence in NTHi, promoting extracellular DNA uptake for nutritional purposes [81,82]. Our study demonstrated that Lung strains utilized isoleucine to the greatest extent compared to ES and PS strains. In contrast, valine was produced only by ES strains.

Our observations of differential metabolic profiling suggest that NTHi strains adapts specifically to the occupied niche. However, based on current knowledge, we are unable to fully explain the underlying reasons for the different metabolic profiles of strains isolated from various body sites. Further in vitro and in vivo studies are necessary to validate the physiological significance of our findings. Nevertheless, our data provide a foundation for the development of targeted site-specific therapeutic approaches for NTHi infections.

Some significant compounds remained unannotated against the available mass spectral libraries under given circumstances of annotating parameters [68,83]. Application of further methods, such as gas chromatography coupled with mass spectrometry should be used for further identification of the significant signals.

The discrepancy between the largely conserved gene content and metabolic pathways, despite low but detectable core-genome SNP variation, and the observed differential metabolic pattern is noteworthy. One possible explanation for this phenomenon is the presence of phase-variable regulons (phasevarions) in human-adapted bacterial pathogens, including NTHi. Phasevarions can lead to differential expression of genes through epigenetic mechanisms, resulting in ON-OFF switching that controls the expression of genes required for pathobiology [19]. Given that core-genome SNP analysis revealed limited genomic divergence (8-116 SNPs) among the strains, the observed metabolic differences are more likely attributable to regulatory or epigenetic mechanisms rather than major gene-content variation. To validate this assumption, further analyses such as transcriptomics and epigenetics profiling would be necessary.

The small and unequal sample size (2 ear smear strains, 2 pharyngeal smear strains, and 5 lower respiratory tract strains) limits the generalizability of our findings to site-specific NTHi strains. Therefore, further investigation with a larger sample size is necessary to confirm our results and define strain diversities in more detail. Additionally, we did not observe sufficient annotated significant metabolic signatures in sample groups related to diagnosis (otitis media and pneumonia) while taking off ’Site’ as covariate control in MetaboAnalyst, and thus did not include them in this study. Also, this study has mainly focused on site specific metabolic variation rather than diagnostic specific variables. As a result, we did not control for diagnostic group as a covariate in MetaboAnalyst when examining the linear relationship among site-specific sample groups. In the metabolomic and statistical sections, we included media control for two main reasons. Firstly, compounds detected equally in the cultivation media could be considered as media components and excluded from statistical analysis. Secondly, compounds from the cultivation medium that were metabolized differentially by different strains could be identified and addressed through statistical analysis.

In summary, this study underscores the significance of metabolomic investigations from a bacterial perspective. A deeper understanding of pathogen-specific metabolic profiles provides insight into survival strategies within the host and informs about site/disease specific microbial communities. Our findings support the hypothesis that NTHi strains adapt to distinct infection sites through metabolic reprogramming, despite possessing a largely homogeneous genomic capacity for metabolite synthesis. The observed metabolic variations suggest that NTHi may modulate nutrient uptake and physiological processes in a site-specific manner. Future research should focus on validating these pathways functionally and assessing their relevance in vivo.

## Materials and methods

### Genome assembly, quality assessment, and annotation

Sequencing was previously performed using Illumina Miseq with 251 bp paired-end reads [20]. Raw reads were quality-checked, trimmed, assembled, and polished using Shovill with SPAdes v.3.14.0 as the assembly algorithm, yielding contig-level assemblies [84]. Assembly completeness was assessed using BUSCO v5.8.2 (Benchmarking Universal Single-Copy Orthologs) with *bacteria_odb10* dataset [85]. Assembly metrics were evaluated with QUAST v5.3.0 [86]. Antimicrobial resistances were identified using ABRicate v1.0.1 with CARD settings against the NCBI AMR database (--db ncbi). Species-level identity was confirmed by FastANI v1.0 via average nucleotide identity comparisons to *H. influenzae* CP000057.2 reference genome [85–87]. The complete genomes of all isolated strains are deposited in the European Nucleotide Archive database (Bioproject; PRJEB43356).

### Functional annotation and comparative genome analysis

Strains exhibiting high completeness (BUSCO scores: 98.2-99.4%) and robust quality assembly (few contigs, high N50) were selected for downstream analysis [21,88]. Functional annotation was performed using both the RAST server (https://rast.nmpdr.org/) and Prokka v1.14.6 to ensure accuracy and completeness [89]. RAST provided subsystem mapping, gene function prediction, and pathway associations. ResFinder 4.6.0 was used to determine anti-microbial resistance genes [90]. Stress adaptation genes (e.g., oxidative stress, heavy metal resistance, osmotic stress) were identified and analysed for potential niche-specific relevance. Core-genome single nucleotide polymorphisms (SNPs) were identified from the core gene alignment generated using Roary, with SNP sites extracted using snp-sites and pairwise SNP distances calculated using snp-dists. The dual annotation (RAST + Prokka) approach minimized gene fragmentation and miss-annotation.

### Pangenome and functional category analysis

Pangenome analysis was conducted using Roary, based on Prokka-generated GFF3 files. Functional categorization of orthologous groups was performed via COG (Clusters of Orthologous Groups) assignment, supporting comparative genomics, PCA, and clustering analyses. COGs enable comparative analysis of gene content across species by grouping genes with shared ancestry and presumed similar functions.

### antiSMASH and KEGG pathway analyses

Biosynthetic gene cluster (BGC) prediction for each of the 9 human pathogenic bacterial strains was performed using antiSMASH v8.0.1, with annotated GenBank (.gbk) files generated via PROKKA as input [91,92]. Processing parameters in antiSMASH were set with detection strictness to strict, and extra features considered for KnownClusterBlast, ClusterBlast, SubClusterBlast, MIBiG cluster comparison, ActiveSiteFinder, RREFinder and TFBS analysis. Comparative BGC profiling was carried out across strains, with selected clusters prioritized for experimental validation of metabolite production. Given the pronounced inter-strain variation in carbohydrate metabolism, functional annotation and metabolic pathway mapping were conducted using the Kyoto Encyclopedia of Genes and Genomes (KEGG) and RAST servers [93–96]. Genes assigned to carbohydrate metabolism were identified, covering CO₂ fixation, central carbohydrate metabolism, polysaccharide and monosaccharide utilization, di-/oligosaccharide metabolism, amino sugars, organic acids, fermentation, one-carbon metabolism, glycoside hydrolases, and sugar alcohol metabolism. Presence/absence patterns were visualized using heatmap analysis. Individual genes from each carbohydrate subpathway were retrieved using KEGG Automatic Annotation Server (KAAS), cross-referenced with PROKKA outputs, and assigned KEGG orthology to determine pathway completeness and differential gene content, revealing potential metabolic adaptations relevant to pathogenicity and host–pathogen interactions [97]. For investigating metabolic potential of detected significant metabolites across the site-specific strains, KAAS was applied with bi-directional best hit method. In KAAS job request for BLAST program, amino acid sequences of the whole genomes were uploaded and search against representative genes of prokaryotes including 8 *Haemophilus* strains (hin, hit, hip, hiq, hif, hiu, hpr and hdu). KEGG orthologies of observed genes for selected pathways and per NTHi strains were retrieved from KEGG Mapper.

### Cultivation and extraction

Each strain of NTHi strains was streaked on chocolate agar (BD) and incubated at 37°C for 24 h. A pure colony grown after 24 h was inoculated into 6 mL chemically defined medium as mentioned by Coleman et al. (2003) and incubated for 24 h at 37°C at an agitation of 160 rpm [98]. The cultivation medium consisted of 194 mL of RPMI 1640 with L-glutamine and phenol red but without 25mM HEPES, 2 mL of a 100 mM MEM sodium pyruvate solution (Sigma-Aldrich), 2 mL of β-NAD^+^ stock, 4 mL of heme-L-histidine stock, 10 mL of a 2 mg/mL uracil solution dissolved in 0.1 M NaOH, and 20 mL of a 20 mg/mL inosine solution prepared by dissolving inosine (Sigma-Aldrich) in deionized water and filter sterilized (0.2 µm pore size). The β-NAD^+^ stock was prepared by dissolving 100 mg of β-NAD^+^ (Sigma-Aldrich) in 100 mL of deionized water and filter sterilized (0.2 µm pore size). The heme-L-histidine stock was prepared by dissolving 0.2 g of L-histidine (Sigma-Aldrich) in 200 mL of deionized water and adding 0.2 g of hemin HCl (Sigma-Aldrich) and 4 mL of 1 M NaOH and steaming over a water bath for 10 min to solubilize the mixture, and filter sterilized (0.2 µm pore size). After measuring optical density at 600nm (OD_600_), one mL of each seed culture was further inoculated into 18 mL of fresh medium (main culture) and incubated for 24 h at 37°C and 160 rpm. Depending on the growth (OD_600_) of different strains, to obtain the OD_600_ > 0.50, the incubation period varied up to 28 h. The main culture of each strain was centrifuged (Avanti J-25 XP) for 30 min at 4°C and 30074 × *g*. The supernatant was filter sterilized (0.2 µm pore size) and stored at -20°C until n-butanol extraction, while the cells were washed with sterile water, liquid N_2_-killed, and stored at -80°C.

For solvent extraction, the supernatant of each culture was mixed with 20 mL n-butanol and shaken vigorously for 20 min. Afterwards, the mixture was centrifuged (Thermo Scientific Sorvall RC 6+ Centrifuge) for 10 min at 10°C and 2290 × *g*. The upper solvent phase was collected, evaporated under a rotary evaporator, redissolved in 1 mL methanol (Millipore), and N_2_-dried to yield the final dry extract ready for LC/MS measurements.

### LC-MS measurement and metabolomics

Samples for LC-MS measurement were prepared at a concentration of 0.1 mg/mL in methanol and analysed by ultrahigh-performance liquid chromatography–electrospray ionization–high-resolution tandem mass spectrometry (UHPLC–ESI–HRMS/MS) using an Elute UHPLC system coupled to a Compact QTOF mass spectrometer (Bruker Daltonics, Bremen, Germany) equipped with a reversed-phase Intensity Solo 2 C18 column (100 mm × 2.1 mm, 2.2 µm; Bruker Daltonics, Bremen, Germany). The UHPLC acquisition method was set with column temperature: 35°C, autosampler temperature: 4°C, sample injection volume: 10 µL with flush volume of 45 µL via partial loop injection. Water with 0.1% formic acid (buffer A) and acetonitrile with 0.1% formic acid (buffer B) were used for mobile phases with the elution gradient of 10% B isocratically for 2 min, then to 100% B in 17 min, washing the column with 100% B for 5 min, and finally recalibration of the column with 10% B for 5 min. The MS acquisition was set up with following parameters; mass spectra peak detection: use maximum intensity, absolute threshold (per 1000 sum.): 25 cts., absolute threshold: 20 cts., peak summation width: 5 pts., source: ESI, end plate offset: 500 V, capillary: 4500 V, nebulizer: 2.2 Bar, dry gas: 10L/min, dry temperature: 220°C, Funnel 1RF and Funnel 2RF: 200 Vpp, isCID energy: 0 eV, hexapole: 50 Vpp, quadrupole Ion energy: 5 eV, collision cell energy: 5 eV, pre pulse storage 5 µs, stepping mode: basic, collision RF: 200-700 Vpp, transfer time: 20-80µs, MS/MS only collision energy: 100-250%, precursor ion cycle time: 0.3 sec, MS/MS absolute threshold (per 1000 sum.): 400 cts., MS/MS absolute threshold: 313, active exclusion: after 3 spectra and 0.3 min, calibration mode: HPC, calibration reference compound: sodium formate, ion polarity: positive, scan mode: MS, scan range: 20-1300 *m/z*. Each sample was measured twice accompanied by external quality control (QC, 0.0125 mg/mL biotin, injection volume of 2 µL) and methanol blank. The raw LC-MS data of 48 samples, including 4 external QC samples, 2 n-butanol blank samples, 4 methanol blank samples, 2 media control samples, and 36 bacterial-culture extracts (encompassing 2 experimental- and 2 technical-replicates) were processed through MetaboScape 2022b Version 9.0.1. In MetaboScape, following parameters were set: minimum number of features for extraction: 4, presence of features in minimum number of analyses: 4, processing method: T-ReX 3D Processing, intensity threshold: 8000 counts, feature signal: area, minimum peak length: 10 spectra, retention time range: 0-22 min and mass range (*m/z*): 50-1300. Furthermore, ion deconvolution was set with EIC correlation: 0.8, primary ion: [M+H]^+^, seed ions: [M+Na]^+^, [M+K]^+^, [M+2H]^2+^ and [M+2Na]^2+^, and common ions: [M+H-H_2_O]^+^ and [M-H_2_O+2H]^2+^.

In MetaboScape, the feature table of metabolites was annotated against different mass spectral libraries including Bruker MetaboBASE Personal Library 3.0 (100679 compounds), MassBank_NIST (21804 compounds), MSMS_Public_Pos_VS17 (21127 compounds) and MoNA (17748 compounds). During annotation against libraries, the parameters were set as *m/z* tolerance of 5 ppm, mSigma value up to 60, and a minimum MS/MS score of 800. The feature table of metabolites was exported as an excel file (.csv) from MetaboScape after filtering out the features eluted before 0.5 min, features lacking MS/MS spectra, features present extensively in n-butanol.

### Statistical analysis

The feature table generated from nine NTHi strains was considered for statistical analysis. It was further cleaned by filtering out all the features present in 4 methanol blank samples. Additionally, features present in only media-control samples were also removed. At the final cleaning step, all those features were kept in the table that were present in at least one set of 4 values representing 2 experimental- and 2 technical-replicates while the rest were completely filtered out from the feature table. Additionally, QC, n-butanol, and methanol blank samples were also removed from the table. In the table, the samples were attributed to four groups: ES, PS, Lung and not applicable (“na”). Lung group consisted of samples from the bacteria isolated from tracheal secretion, bronchial secretion and bronchoalveolar lavage. Media control sample was assigned to “na” group.

The final feature table was split into data table consisting of sample names versus feature names with their peak area and a metadata table consisting of two columns: sample names and sampling sites (ES, PS, Lung, na). In the data table, samples were kept in rows and features were kept in columns.

In MetaboAnalyst 6.0, statistical analysis module is used for the comparative analysis of metabolites produced by NTHi strains isolated from different body sites [99–101]. The data and metadata files were uploaded by considering peak intensities as data type. Data filtering step was skipped while manual data cleaning was done before processing data in MetaboAnalyst. Afterwards, the data were normalized by log transformation (base 10). Finally, principal component analysis (PCA) and linear model were performed by considering each group as reference and the other group as contrast with FDR-adjusted *p*-value cutoff of 0.05 without covariate adjustment. The significant metabolites were tabulated with their logFC, *p*-value, and adj *p*-value. Box plots of each significant metabolite in different comparison groups were generated using the *tidyverse* package in R (version 4.5.1).

## Acknowledgements

Not applicable

## Data availability statement

The data that support the findings of this study are openly available in MassIVE at doi:10.25345/C58S4K27R, reference number MassIVE MSV000099632.

## Supporting information

**S1 Figure Whole genome sequence-based phylogenetic tree of 172 *H. influenzae* isolates derived from Diricks et al.(2022).** The approximately maximum-likelihood tree was previously constructed based on 104 core genes using FastTree. Local bootstrap values below 0.9 are highlighted with blue circles. NTHi clades were previously determined using the bioinformatic tool patho_typing based on gene absence/presence patterns. For isolates that were not classified by patho typing in one out of six previously defined clades (TND), clade affiliation was inferred from phylogenetic positioning. Clade VI was further divided into subgroups a and b based on their positions within the current tree.

**S2 Figure GBDP distance-based phylogenetic tree obtained from 16S rRNA gene sequences generated using FastME 2.1.6.1.** Using the GBDP distance formula d5, branch lengths are proportionate to distances. Bootstrap support values >60% from 100 pseudo-replicates are indicated above branches, with a mean branch support of 63.0%. The tree was rooted at the midpoint. Species with blue asterisks (*) are the strains currently studied.

**S3 Figure MS^2^ spectral match of the metabolites from 9 NTHi isolates against MS^2^-libraries.**

**S4 Figure 3D PCA plot of site-specific samples based on metabolites intensities with metadata overlay.** (ES = ear sample, PS = pharyngeal sample, na = media control, OM = otitis media and P = pneumonia).

**S1 Table Genomic assembly quality, structural features, and functional landscape of *Haemophilus influenzae* strains.**

**S2 Table Comparative genomic visualization of NTHi strains generated using Proksee.**

**S3 Table Genome-wide phylogenomic analysis of Haemophilus influenzae isolates based on GBDP clustering.**

**S4 Table Distribution of KEGG orthologs (KOs) and corresponding gene annotations linked to carbohydrate metabolism**. The listed KOs and gene annotations encompass major carbohydrate utilization and biosynthetic pathways across *Haemophilus* strains. The table highlights strain-specific presence/absence patterns reflecting metabolic conservation and diversity within the carbohydrate metabolic network.

**S5 Table Biosynthetic gene cluster (BGC) regions of different NTHi strains analysed by antiSMASH version 8.0.2.**

**S6 Table List of 38 metabolites detected using LC-MS and their signal intensities in the respective samples.**

**S7 Table Significant metabolites with P-cutoff: 0.05 (FDR) varied in different groups without covariate adjustment.** The compared groups are ES vs Lung (a), ES vs PS (b) and Lung vs PS (c).

**S8 Table Annotated KEGG orthology groups for different pathways present in individual bacterial isolates.**

